# GLnexus: joint variant calling for large cohort sequencing

**DOI:** 10.1101/343970

**Authors:** Michael F. Lin, Ohad Rodeh, John Penn, Xiaodong Bai, Jeffrey G. Reid, Olga Krasheninina, William J. Salerno

## Abstract

As ever-larger cohorts of human genomes are collected in pursuit of genotype/phenotype associations, sequencing informatics must scale up to yield complete and accurate genotypes from vast raw datasets. Joint variant calling, a data processing step entailing simultaneous analysis of all participants sequenced, exhibits this scaling challenge acutely. We present *GLnexus* (GL, Genotype Likelihood), a system for joint variant calling designed to scale up to the largest foreseeable human cohorts. GLnexus combines scalable joint calling algorithms with a persistent database that grows efficiently as additional participants are sequenced. We validate GLnexus using 50,000 exomes to show it produces comparable or better results than existing methods, at a fraction of the computational cost with better scaling. We provide a standalone open-source version of GLnexus and a DNAnexus cloud-native deployment supporting very large projects, which has been employed for cohorts of >240,000 exomes and >22,000 whole-genomes.

## 1. INTRODUCTION

Sequencing large cohorts of human genomes to associate germline variation with phenotypes and disease risk is a mainstay of modern genome science. Variant calling methodologies for short-read sequencing readily ascertain single-nucleotide variants (SNVs) and small insertions/deletions (indels) relative to the reference genome assembly, typically recorded in Variant Call Format (VCF) [1]. In addition to describing variants for one sequenced participants, VCF can present genotypes for an entire cohort, in a 2-D matrix of variant sites and study participants, filled with the diploid genotypes and quality-control (QC) measures – referred to as a Project VCF (pVCF). The pVCF’s matrix format facilitates downstream calculation of allele frequencies and association testing statistics in the cohort or any phenotype-defined subsets of it.

The 1000 Genomes Project and other early cohort sequencing efforts developed methodologies for pVCF production based on pooled analysis of all available sequence read mappings across the cohort [2–8], maximizing the evidence used to resolve noisy genotypes, but creating formidable data processing challenges as cohort sizes and sequencing depths both increased. To ameliorate this, intermediate formats were introduced to compactly summarize key features of potential variants and genotypes from each participant’s read mappings separately, which can be reprocessed jointly into pVCF or a similar matrix format [9]. Among the most successful of these has been an adaptation of Genome VCF (gVCF) – a VCF extension supplementing variant sites with coverage information to distinguish reference-equivalent from uncertain regions [10] – used for pVCF production in the Genome Analysis Toolkit (GATK) [11,12].

Joint variant calling, the process of producing the pVCF matrix from the set of gVCFs or equivalents, has several challenges which increasing cohort sizes tend to exacerbate, pressing for continued methodological innovation to keep pace. (1) *variant representation*: large cohorts frequently exhibit multiple and overlapping alleles at a given genome locus, requiring the pVCF to trade off between completeness and mutual-exclusivity. Compounding this, VCF admits multiple ways to write the same variant, necessitating algorithms to unify them [13,14]. (2) *joint genotyping*: the initial gVCF genotype calls for each participant, whether variant or reference-equivalent, can be refined in light of the cohort-wide allele frequencies and error patterns [3,15,16]. (3) *scalable data processing*: while each gVCF is relatively compact, pVCF production still involves reprocessing the entire cohort at once, with super-linear scaling in runtime and output size as the cohort grows and more rare variants are genotyped.

This article presents *GLnexus* (GL, Genotype Likelihood), cloud-based software for joint variant calling designed to scale up to any cohort size foreseeable under the gVCF/pVCF data model. GLnexus combines scalable joint calling algorithms with a persistent database for querying the gVCF data corpus that grows efficiently as additional participants are sequenced. We evaluate GLnexus pVCF quality in a cohort of 50,000 exomes and describe its application on >240,000 exomes and >22,000 whole genomes. GLnexus’ algorithms are available in a standalone, open-source tool, while a DNAnexus cloud-based framework facilitates deployment for large-scale data production.

## 2. ALGORITHM & SYSTEM DESIGN

### 2.1 Joint variant calling in large cohorts

To frame GLnexus’ design, we begin with a general definition of gVCF-to-pVCF joint variant calling, independent of the specific implementation. The gVCF file representing a study participant decomposes the length of each reference chromosome (contig) into a set of candidate variant sites and intervening stretches without evident variation. Each such record encompasses one or more reference base positions and specifies a set of candidate alleles replacing the reference subsequence in the participant’s genome, beginning with the reference allele itself, followed by zero or more alternate alleles, and lastly a catch-all entry symbolizing “other.” These *k* alleles imply *k*(*k*+1)/2 candidate diploid genotypes. Each record includes a vector of genotype likelihoods Pr(*D* | *G* = *g*), the probability of the observed data (alignments of sequence reads to the locus) assuming the specific genotype *g*, according to a probabilistic model intrinsic to the gVCF caller. The maximum-likelihood genotype call, argmax*_g_* Pr(*D* | *G* = *g*), and various QC measures are also reported; in stretches without apparent variation, this is reference-homozygous with QC measures reflecting read coverage and quality.

Joint calling a set of gVCFs to pVCF involves, first, deriving a set of “unified” variant sites representing all discovered alleles passing QC thresholds (**Figure 1**). Then, for each participant, the pertinent gVCF alleles, genotypes, likelihoods, and QC measures are “projected” onto the unified sites, including reference-homozygous calls where applicable, sometimes using multiple gVCF records to inform one pVCF matrix entry and vice-versa. Lastly, the maximum likelihood genotype calls in each gVCF may be revised to maximum *a posteriori* calls, argmax*_g_* Pr(*G* = *g*) Pr(*D* | *G* = *g*), where the prior might factor in (1) *de novo* terms from population genetics theory, (2) site-specific empirical estimates of allele frequencies and error patterns, and (3) participant-specific terms reflecting ethnicity, relatedness to other participants, or experimental batch.

**Figure 1.**
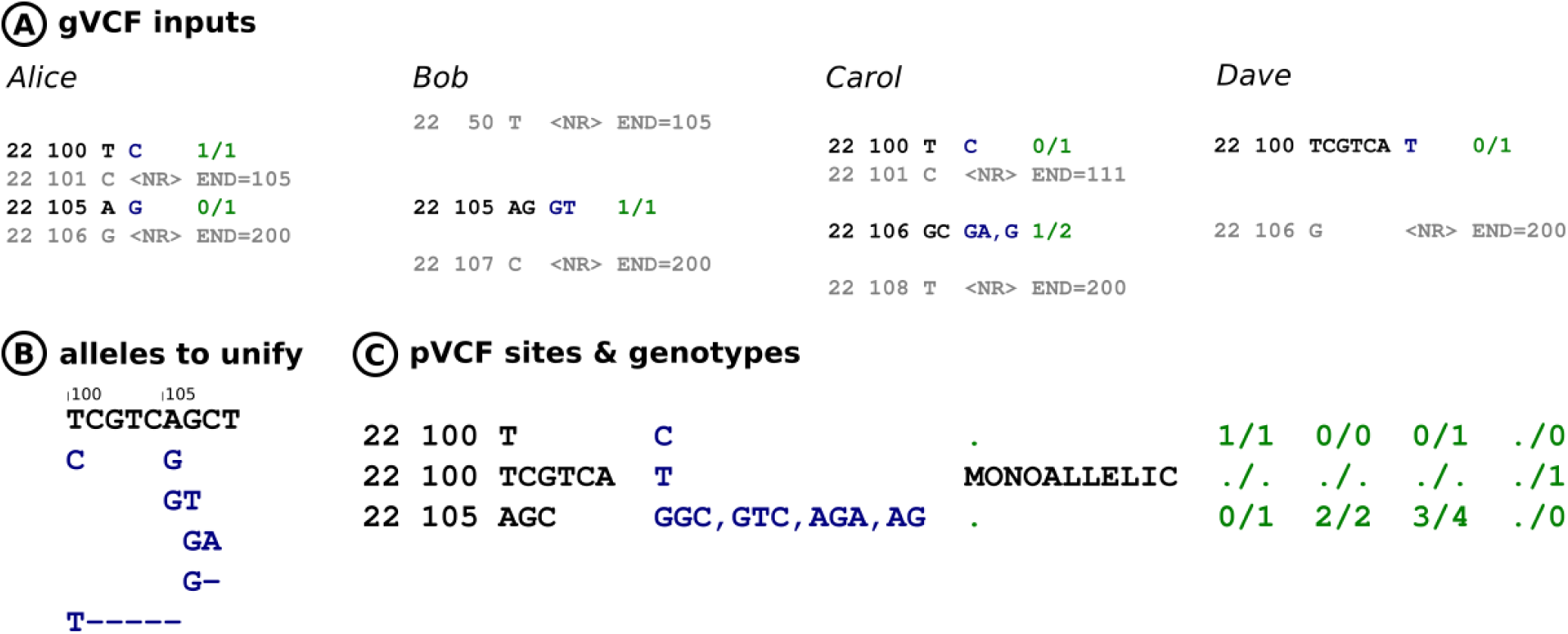
Allele unification in joint variant calling. (A) Example abbreviated gVCF records for four participants, giving genome position, reference and alternate alleles, and initial called genotypes. Gray records indicate sequencing coverage for regions with no apparent variation. (B) Schematic view of the alternate alleles seen across the four gVCF inputs; they cluster into two sites except for a spanning deletion allele. (C) Example pVCF representation for these variants, with two multiallelic sites and a third “monoallelic” site representing the deletion allele which could not be unified into the multiallelic sites without introducing phase constraints artificially. The input alleles, genotype calls and (not shown) QC measures from the input gVCF records must be “projected” onto the pVCF site representation.

With this framework, we discuss selected features of GLnexus’ joint-calling implementation that proved significant for use with very large cohorts.

#### Hybrid allelic representation

To facilitate downstream summary statistics without double-counting, ideal unified sites would be completely non-overlapping, with mutually-exclusive alleles. But, this can be infeasible when deletion and multi-nucleotide variants span several other, generally unphased alleles. Such complications are unusual within any one genome but arise more frequently amongst the many rare alleles inevitably found in larger cohorts.

GLnexus’ allele unification algorithm considers all discovered alleles (meeting QC thresholds) in descending order of frequency. It greedily unifies these into non-overlapping multi-allelic variant sites, padding alleles with reference bases and recognizing synonymous representations as needed. Upon encountering an allele that doesn’t unify into the multi-allelic sites constructed so far, it generates a specialized “mono-allelic” site, conveying for each participant the presence (copy number) of that allele *only* [17]. The other overlapping sites convey the copy numbers of the reference and any other alleles, with missing or partial genotype calls for carriers of the separated allele.

This hybrid representation (**Figure 1C**) includes in the pVCF all discovered alleles passing QC, usually within non-overlapping sites; while ensuring that, for any given reference base position, it indicates presence of at most two allele copies in each participant genome. Its trade-offs include presenting two distinct kinds of pVCF sites and complicating simple alleles when lengthier ones overlap them. Alternatives, seen with other methods, are to omit difficult-to-unify alleles and/or to generate more overlapping sites, possibly with redundant calls.

#### Empirical frequency prior

Population allele frequencies can improve genotype calls in the pVCF matrix. For example, upon encountering a locus covered only by a few reads all exhibiting a certain alternate allele, a gVCF caller would reasonably report a homozygous-alternate genotype (with poor QC measures). With further knowledge that this allele is very rare, joint calling can statistically “shrink” the called genotype to heterozygous, since a higher threshold of evidence is needed to justify the homozygous call. Borderline heterozygous calls might similarly revise to major-allele homozygous. With deep sequencing coverage, such poorly-covered regions arise infrequently in any one participant genome; yet they remain an inevitable source of noise throughout the pVCF genotype matrix for the whole cohort.

Initially, we formulated a genotype prior with Hardy-Weinberg equilibrium genotype frequencies implied by the empirical allele frequencies estimated from the gVCF cohort. Testing this on cohorts of tens of thousands, we noted an undesirable effect also observed with other joint-calling methods [18]: sensitivity to very rare alleles (singletons or private to a family) declined with increasing cohort size *N*, owing to their declining estimated allele frequencies proportional to 1/2*N*. We therefore reformulated the prior with a ramp function on the estimated allele frequency; intuitively, this stops escalating the evidence threshold to call rare genotypes once the cohort is large enough to confidently distinguish rare and common alleles (a few thousand participants). This modified prior ensures consistent sensitivity to rare alleles as the available cohort grows.

### 2.2 gVCF database for incremental reprocessing

To produce a pVCF, GLnexus must visit each unified site and process pertinent records from all cohort gVCFs — a challenging data access pattern, given that the records are variable-length and sometimes overlapping, and that the compressed gVCFs total many terabytes for modern cohorts, in one file per participant. Furthermore, to support sequencing projects carrying on over months and years, we sought to streamline delivery of successively larger pVCFs, minimizing repeated processing of the inputs.

GLnexus imports gVCF files into a persistent database backed by RocksDB (http://rocksdb.org), a key/value storage library providing efficient scans of ordered key ranges, parallel bulk load, storage compression, and online defragmentation. During import, each gVCF record is assigned to a 30-kilobase reference genomic range “bin” according to its position. The records in each bin are serialized to a binary message and stored in RocksDB using the key <bin genomic range, participant> (**Figure 2A**). To retrieve records near a certain genomic range, GLnexus asks RocksDB for an ordered scan of keys beginning with the relevant bin(s), thus efficiently “slicing” the cohort gVCF corpus. A binary search index stored alongside each bin further expedites locating the gVCF records which directly overlap the query range. (Records overlapping multiple bins are repeated in each bin, and the query logic hides duplication while scanning multiple bins.)

**Figure 2.**
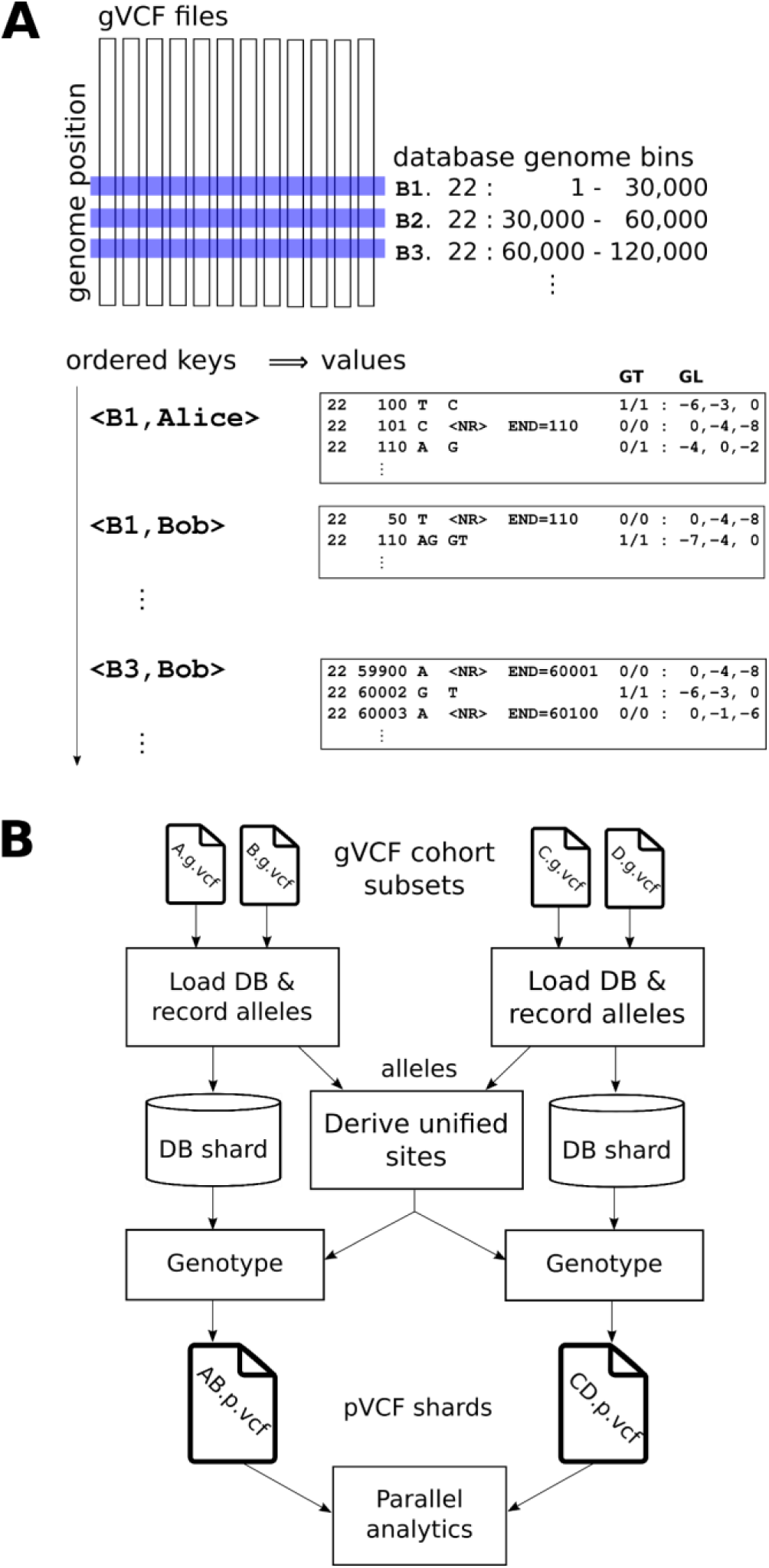
GLnexus database and workflow. (A) To expedite access by genome position, GLnexus loads the gVCF corpus into an ordered key-value database supporting efficient scans of contiguous key ranges. Records from each gVCF file are assigned to a 30-kilobase genome position bin. Bins from each participant are keyed so that all records overlapping a given genome position can be retrieved with a database key range scan. The database values include gVCF records in htslib’s binary format, including genotype likelihoods, and a position search index. (B) Distributed joint-calling cloud workflow, illustrated with two “shards” of the cohort. gVCF files are loaded into a database using separate compute nodes for each shard. Alleles are simultaneously recorded and sent to a central process for unification into pVCF sites. These sites are genotyped across the cohort shards by revisiting records stored in the gVCF database, producing pVCF shards with corresponding rows, suitable for loading into a parallel analytics environment such as Apache Hadoop or Spark.

When additional participants’ gVCF files subsequently become available, the database can incorporate the new bins throughout the ordered key space without immediately rewriting existing stored data, owing to RocksDB’s log-structured merge-tree scheme [19]. pVCF production can then proceed on the full cohort or, by taking advantage of the database’s efficient random lookup, on any subset of participants or sites.

### 2.3 Parallel joint calling workflow

GLnexus employs a three-stage computational workflow to generate pVCF starting from gVCF files. (1) Loading the files into the database. During this process, the system records the candidate alleles seen along with their QC measures. (2) The discovered alleles are QC-filtered and unified into variant sites. (3) The unified sites are genotyped across the cohort by revisiting the gVCF data previously stored in the database, generating pVCF. Internally, GLnexus parallelizes each stage — loading by file, unification by chromosome, and genotyping by site — utilizing up to dozens of threads on one compute node efficiently.

Furthermore, the operations can be distributed across many compute nodes by “sharding” the cohort (**Figure 2B**). In this more complex deployment, each node loads a subset of the gVCF files, transmits the discovered alleles for centralized unification, receives back the sites for genotyping, and finally generates pVCF. The resulting pVCF shards — each presenting the exact same sites and alleles for a subset of the cohort — are suitable for use as a distributed dataset in a parallel analysis environment such as Apache Hadoop or Spark, which can also be used to materialize a monolithic pVCF file by column-binding the shards, if so desired.

Our cohort-sharding strategy has three notable advantages over approaches that divide up the genome for separate processing. (1) It more naturally accommodates incremental growth of the gVCF cohort as additional participants are sequenced. (2) It is less necessary to cut the genome into ever-smaller ranges, which can cause complications with variants near the cut points. (3) It is less prone to outlier running times among the several compute jobs owing to genome regions with unusually dense multiallelic variation, which tends to worsen with cohort size.

## 3. VALIDATION STUDY WITH 50,000 EXOMES

To benchmark GLnexus, we reanalyzed *N*=50K deep whole-exome sequences from the DiscovEHR cohort, population genetics of which have been published previously [20]. We generated gVCFs for these exomes using GATK HaplotypeCaller 3.8 and joint-called the exome capture targets on chromosome 2 (2.6 Mbp covered) using both GATK (CombineGVCFs in batches of *N*=500, followed by GenotypeGVCFs) and GLnexus. Low-complexity regions were then excluded from comparisons. With default settings, the two methods produced generally similar pVCFs, as expected given identical gVCF inputs from deep sequencing, with roughly 280,000 distinct alternate alleles, ~42% singleton (**Table 1**). At corresponding sites, the pVCF matrices showed discordant genotype calls (variation indicated in one but not the other) in only 0.00073% of 12.8 billion cells and 0.12% of 80 million cells with any variation in either pVCF.

**Table 1.** Chromosome 2 pVCF characteristics for *N*=50,000 DiscovEHR exomes

|  | GLnexus | GATK3.8 |
| --- | --- | --- |
| pVCF sites (chromosome 2) | 268 580 | 259 819 |
| Unique (no overlap in other) | 10 851 | 92 |
| With >1 alternate alleles | 15 849 | 11 618 |
| Overlapping | 4 017 | 7 381 |
| Alternate alleles | 287 363 | 272 081 |
| SNV | 265 262 | 255 407 |
| indel | 22 101 | 16 674 |
| Alternate alleles, $\geq 0.1\%$ freq | 9 857 | 9 877 |
| SNV | 9 498 | 9 518 |
| indel | 359 | 359 |
| Alternate alleles, singleton | 121 885 | 117 883 |
| SNV | 111 510 | 109 824 |
| Indel | 10 375 | 8 009 |

Beneath this overall concordance, GLnexus included more alleles in the pVCF sites — especially rare indels, of which GLnexus carried over 33% more (**Figure 3A**). Of 3,228 biallelic indel sites in GLnexus’ pVCF with no overlap in GATK’s pVCF, nearly all (98.7%) had ten or fewer copies called in the cohort; 45.2% were singletons, with median Genotype Quality (GQ) score of 74 (Phred scale), and the remainder were called in multiple participants with median quality 57.

**Figure 3.**
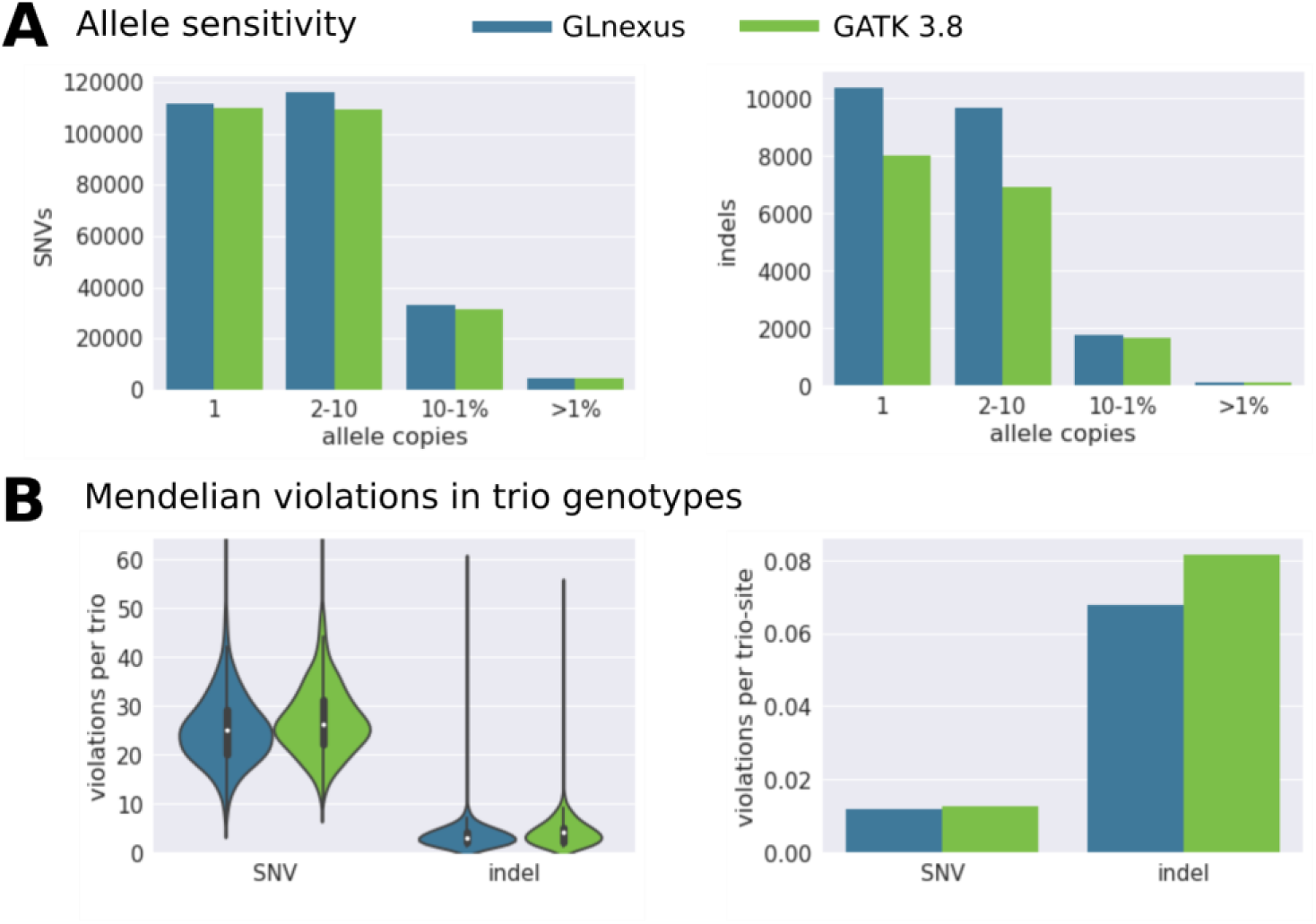
Sensitivity and error rate in GLnexus and GATK pVCFs for 50,000 chromosome 2 exomes. (A) Number of distinct alternate alleles in pVCF sites, decomposed by copy number in called genotypes. (B) Violations of Mendelian inheritance indicated within genotype calls for 877 trios in the cohort. Left, distribution of violation count per trio on chromosome 2. Right, rate of violations per trio & site exhibiting SNV or indel variation. The GLnexus pVCF exhibits both higher sensitivity and lower Mendelian violation rate, especially with indels.

Since both methods have configurable thresholds, we sought to verify that GLnexus didn’t merely sacrifice quality for sensitivity relative to GATK. Focusing on the 877 child, mother, and father trios within the 50K cohort, we counted apparent Mendelian inheritance violations in their called genotypes; i.e. sites where the two allele copies called in the child lacked copies called in each parent. We used these as a proxy for genotype errors, since far more violations were indicated than a plausible number of *de novo* mutations per child. (Joint calling can mitigate only a fraction of such errors, which have numerous sources throughout the sequencing and bioinformatics.) The GLnexus pVCF tended to indicate fewer Mendelian violations, and fewer violations per site with variation (**Figure 3B**). This difference was most pronounced for indels: the GATK pVCF indicated a Mendelian violation in 8.2% of 44,107 genotype triplets with indel variation, reflecting the challenges indels still present for short-read sequencing, while GLnexus’ corresponding rate was 6.8%.

We also compared unity of variant representation. The GLnexus pVCF included a higher proportion of sites with multiple alternate alleles (5.9% vs. 4.5%), and a smaller proportion of sites overlapping one or more others (1.5% vs. 2.8%); these differences were mostly due to GLnexus’ inclusion of deletions at multiallelic sites, when possible. Monoallelic sites, generated when multiallelic unification was not possible, comprised 0.35% of sites in the GLnexus pVCF, overlapping 1.1% of the primary sites.

### Joint calling scaling from *N*=1K to *N*=50K

Next, we joint-called random subsets of the 50K cohort numbering 1K, 5K, 10K, and 25K participants, each nested within the larger sets, to simulate how pVCF properties would evolve as a sequenced cohort grows. As expected, the number of sites, proportion of sites with multiple alternate alleles, and proportion of overlapping sites each grew steadily with *N* (**Table 2**). The proportion of sites with only rare alleles (total alternate frequency <0.1%) climbed from 68% up to 96%, while at the same time the proportion of singleton biallelic sites declined from 58% down to 41%, as the larger cohorts tended to capture more carriers of each rare allele. The corresponding GATK pVCFs showed proportionate trends.

**Table 2.** GLnexus chromosome 2 pVCF sites with nested subsets of the 50,000 DiscovEHR exomes

|  | *N*=1K | *N*=5K | *N*=10K | *N*=25K | *N*=50K |
| --- | --- | --- | --- | --- | --- |
| pVCF sites (chromosome 2) | 29 179 | 76 928 | 114 922 | 188 336 | 268 580 |
| With >1 alternate alleles | 276 (.95%) | 1 539 (2.0%) | 3 073 (2.7%) | 7 842 (4.2%) | 15 849 (5.9%) |
| Overlapping | 69 (.24%) | 349 (.45%) | 627 (.55%) | 1 784 (.95%) | 4 017 (1.5%) |
| With <0.1% alternate freq | 19 845 (68%) | 67 224 (87%) | 105 113 (91%) | 178 487 (95%) | 258 722 (96%) |
| With 1 singleton alternate | 16 811 (58%) | 43 534 (57%) | 61 088 (53%) | 86 934 (46%) | 109 015 (41%) |

To illustrate our modified population frequency prior, we checked how many of the sites in the *N*=5K pVCF had no overlapping site in the superset 10K, 25K, and 50KpVCFs. Using GATK, this number climbed from 309 of the 77,138 5K sites (0.40%) absent in the 10K pVCF, to 465 (0.60%) absent with 25K and 492 (0.64%) absent with 50K. In contrast, GLnexus lost sensitivity only to a residual number (fewer than 10) of the sites in its 5K pVCF with larger cohorts, which were artifacts of filtering low-complexity regions.

The sites absent in the larger GATK pVCFs tended to represent very rare alleles with below-average call quality (median GQ 71 among the 492 sites absent in the 50K pVCF), often heterozygous with multiple, but far below 50%, supporting reads for the alternate allele – perhaps reflecting segmental duplications or other difficult-to-map regions. The genotype quality at incrementally lost sites tended to increase with each larger cohort, consistent with our observations designing GLnexus’ frequency prior.

The pVCF output from joint-calling would typically undergo further QC filtering suited to specific downstream analyses, such as identifying candidate *de novo* mutations. Our observations show that GLnexus produces a similar but more comprehensive, accurate, and unified pVCF substrate for downstream analysis.

### Runtime comparison

Lastly, we tallied the computational core-hours consumed to joint-call the 1K, 5K, 10K, 25K, and 50K nested cohorts (**Figure 4**). GLnexus incurred a fraction of the compute time compared to GATK in each case, and the relative difference magnified as the cohort grew, from 3.8-fold fewer core-hours for 1K up to 6.6-fold fewer for 50K. We made these runtime comparisons conservatively, by starting each GLnexus run from the original gVCF files, and including a final Spark-based stage to generate a single pVCF file. We’d therefore expect GLnexus’ relative costs to be lower in routine operations, growing the cohort incrementally and delivering sharded pVCF to the downstream environment.

**Figure 4.**
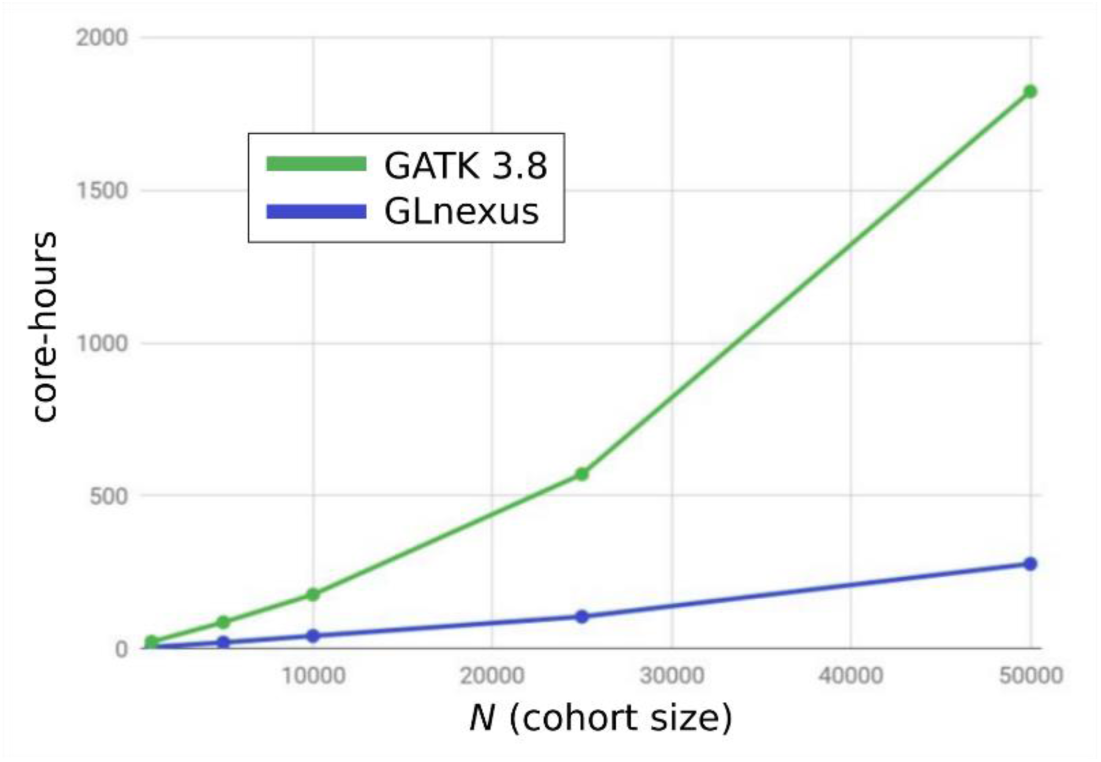
Runtime scaling for joint calling chromosome 2 exomes using GATK and GLnexus Nested sets of *N*=1K, 5K, 10K, 25K, and 50K participants were tested, starting from original gVCF files in each case. GLnexus uses a fraction of the compute time and shows more favorable scaling as the cohort grows.

Contributing factors to GLnexus’ relative efficiency include its gVCF-optimized database, performant C/C++ coding, and different empirical prior estimators. On the other hand, GLnexus is designed for diploid genotyping, in contrast to GATK’s model supporting higher ploidies, and does not compute all the same QC measures.

GATK version 4.0 recently introduced new tools to scale up joint calling. Our initial attempts have not yet observed them to operate more efficiently overall than GATK 3.8 with these cohorts. We plan to update these observations with continued efforts.

## 4. AVAILABILITY

We provide a standalone, open-source version of GLnexus at https://github.com/dnanexus-rnd/GLnexus. It includes the joint calling algorithms in a command-line tool that processes gVCF input files to a pVCF output file. This multithreaded tool can utilize a powerful server with dozens of threads, while the external database lets it operate on datasets larger than the available random-access memory. The DNAnexus cloud-native version of GLnexus supports extremely large cohorts by scaling out on many compute nodes, incremental gVCF database growth and reprocessing, and pVCF delivery as a distributed dataset rather than a monolithic file. The two versions produce identical scientific results, differing in these scalability and production-oriented features.

GLnexus has a declarative configuration scheme enabling it to interpret gVCF inputs from several upstream variant calling tools and accommodate others in the future. In addition to GATK HaplotypeCaller used above, suitable configurations are included for Platypus [7], xAtlas [21], DeepVariant [22], Sentieon DNAseq [23], and Edico DRAGEN, though not all have been calibrated to the same degree. Additionally, GLnexus can be operated in a simplified mode to merge input gVCFs but skip QC filtering and genotype recalculation.

## 5. APPLICATIONS

The Regeneron Genetics Center deploys GLnexus for joint calling on several exome sequencing cohorts and cohort unions. The largest to date encompassed 243,953 participants, processing 33 Terabytes (TB) of compressed gVCF inputs to 6.8TB of compressed pVCF, with 2.7 trillion genotype matrix entries in the 39Mbp exome capture targets. The distributed GLnexus workflow delivered this in approximately 36 hours wall time using 1,600 threads, roughly 14 thread-minutes per exome.

The Human Genome Sequencing Center at Baylor College of Medicine used the distributed GLnexus workflow to merge xAtlas gVCFs [21] generated from 22,609 whole genomes (36.6x median coverage). The combined CRAM-to-pVCF workflow generated a 6.7 trillion genotype matrix from 481 TB of CRAM files in approximately 35 hours wall time and less than 7 core-hours per WGS sample. To minimize batch effects within this cohort sequenced across several NIH-sponsored genome centers, the CRAM files were all generated using the “functional equivalence” protocol for WGS raw data processing [24]. The standalone version of GLnexus has also been used to merge xAtlas gVCFs for 16,521 exomes in 30 wall hours on a 32-thread compute node.

## 6. DISCUSSION

Joint-calling with the cohort sharded across compute nodes will enable GLnexus to scale up to any *N* foreseeable with the gVCF/pVCF data model for short-read sequencing. Eventually, long- or linked-read sequencing of large cohorts may motivate a new paradigm focusing on lengthy haplotypes over isolated variant sites, demanding new algorithms and tools; but GLnexus’ architecture is well-suited to serve current and approaching needs.

The pVCF’s dense genotype matrix is notably space-inefficient for large cohorts: with *N*=50K, 96% of sites had total alternate allele frequency <0.1% (**Table 2**), implying that the vast majority of the matrix consists of reference-homozygous or non-called entries (and their QC metrics, typically more space-consuming than the discrete genotypes). Numerous novel formats and advanced data structures have been proposed recently to exploit this sparsity for impressive efficiency gains. In view of VCF’s wide adoption for interoperable data exchange, however, there may also be a role for an incrementally evolved “sparse VCF,” standardized through a community forum such as the Global Alliance for Genomics and Health.

Genome-wide association studies of multiple separate cohorts can be aggregated through meta-analysis of summary statistics, with much less risk to the participants’ genetic privacy compared to sharing full genotypes [25,26]. This increases discovery power through larger *N* than otherwise feasible owing to cost, recruitment challenges, and data-sharing policy restrictions. These meta-analysis techniques, which rely on genotype data summaries such as site covariance to control potential confounders, may be challenging to fully extend from microarray- to sequencing-based studies, which inherently ascertain different sites in different cohorts, among other distinctive batch effects. GLnexus’ sharding scheme harmonizes the representation of all discovered variants through exchange of similar summary information between cohort subsets on different compute nodes. Although presently implemented to scale within one datacenter, we plan to extend this to harmonize sequenced cohorts held in separate repositories, without centralizing individual-level genotypes, to facilitate federated association meta-analysis.

## 7. ACKNOWLEDGEMENTS

We thank Yifei Men for code contributions; Yih-Chii Hwang, Brett Hannigan, Andrew Carroll, Jesse Farek, Adam Mansfield, and Alicia Hawes for analysis support; and Angela Anderson, George Asimenos, Peter Nguyen, and Brad Chapman for drafting advice.

xAtlas/GLnexus applications are supported by the NHGRI Centers for Common Disease Genomics (5UM1HG008898-02).

